# Cellular mechanism linking endoplasmic reticulum inheritance and cell cycle regulation of the nuclear genome

**DOI:** 10.1101/2025.09.04.674221

**Authors:** Ya-Shiuan Lai, Jesse Chao, Maho Niwa

## Abstract

Endoplasmic reticulum (ER) stress triggers activation of the ER surveillance (ERSU) pathway— a critical protective mechanism that transiently halts cortical ER inheritance to daughter cells and arrests cytokinesis by septin ring subunit Shs1 re-localization to the bud scar in response to ER stress. Once ER functional homeostasis is re-established, cells resume normal cell cycle progression; however, the molecular circuitry linking ER integrity to cell cycle regulation has remained largely unresolved. Here, we show that ER stress selectively disperse Bud2, a GAP for Bud1/Rsr1, severing its canonical role in cell polarity while integrating it into ER homeostasis signaling. Bud2 dispersion results in accelerated spindle pole body (SPB) duplication, spindle misorientation, defects in nuclear migration, and genome segregation errors under ER stress. Strikingly, a C-terminal truncation of Shs1 (*shs1-ΔCTD*) recapitulated the ER stress–induced dispersion of Bud2 phenotype even in the absence of ER stress, and delayed cell-cycle re-entry after ER homeostasis was regained—despite normal occurrence of typical ERSU hallmark events. Notably, Bud2 overexpression rescued the growth defects of *shs1-ΔCTD* mutants after ER homeostasis was re-established. Collectively, our findings reveal a new mechanistic axis whereby ER integrity coordinates organelle inheritance, cytoskeletal organization, and nuclear division via selective control of Bud2 and Shs1, establishing a direct regulatory bridge between ER status and mitotic fidelity.

## Introduction

The faithful inheritance of cellular components—including an error-free genome and functional organelles—is essential to ensure that daughter cells are fully equipped for independent function. In the model organism *Saccharomyces cerevisiae* (yeast), our previous work identified a cell cycle checkpoint that is triggered when endoplasmic reticulum (ER) homeostasis is compromised, resulting in cell cycle arrest at cytokinesis^1, 2^. We termed this checkpoint the ER surveillance (ERSU) pathway. By identifying both the molecular players and the underlying mechanism, we characterized a unique checkpoint that operates independently of the unfolded protein response (UPR)—the canonical signaling pathway that maintains ER homeostasis^3, 4^. Specifically, the ER-membrane proteins Rtn1 and Yop1 detect perturbations in ER function, leading to activation of the MAP kinase Slt2^5, 6^. These events initiate ERSU-specific responses, including blocked ER inheritance, mislocalization of the septin ring, and a temporary halt in cytokinesis^1, 7-9^.

Because cell division is arrested at cytokinesis, ERSU induction must be tightly coordinated with cell cycle progression to safeguard proper chromosome segregation, although the molecular details remain unknown. Importantly, the arrest is reversible: once ER homeostasis is restored through UPR activation, cells resume the cell cycle. This raises compelling questions about how ER-derived signals are integrated with nuclear cell cycle events, and how the ER stress-induced cytokinesis block is coordinated to permit timely re-entry into the cell cycle.

To address these questions, we investigated how blocking ER inheritance during ER stress coordinates with nuclear events linked to genome duplication and segregation. In *S. cerevisiae*, septin subunits assemble into a ring at the site of polarized growth, known as the incipient bud site, which is defined by the accumulation of the small GTPase Cdc42 and its upstream effectors^10,11^. Our earlier studies showed that under ER stress, the Shs1 septin subunit relocates to the bud scar or cytokinetic remnants (CRMs). This re-localization coincides with loss of Cdc42 from the incipient bud site, preventing bud emergence in ER-stressed wild-type (WT) cells^8^.

Strikingly, we found that under ER stress, Cdc42 becomes mis-localized to the bud scar but is rapidly inactivated^8^. Under normal conditions, Cdc42 accumulation at the bud scar is prevented by Nba1^12^. Nba1 remains at the bud scar during ER stress; nonetheless, Cdc42, along with its guanine nucleotide exchange factor (GEF) Cdc24 and the Bud1/Rsr1 GTPase, also accumulates there. This suggests that Cdc42 at the bud scar is rapidly inactivated even in the presence of Cdc24 and Bud1/Rsr1^8^. The molecular mechanism underlying rapid Cdc42 inactivation at the bud scar during ER stress, however, remains unclear.

Although the ER stress-induced cytokinesis block is reversible, the timing of re-entry into the cell cycle can vary. We observed that cells expressing a C-terminally truncated version of Shs1 (*shs1-ΔCTD*) exhibit a pronounced delay in cell cycle re-entry after restoration of ER homeostasis. Notably, *shs1-ΔCTD* cells, like WT cells, display both ER inheritance and cytokinesis blocks during ER stress, indicating that removal of Shs1 from the septin ring at the bud neck—rather than its mislocalization to the bud scar—is sufficient to trigger ERSU events. However, the delayed recovery in *shs1-ΔCTD* cells suggests that mislocalized Shs1 at the bud scar plays a specific regulatory role in coordinating cytokinesis blockade with orderly cell cycle re-entry, rather than simply halting division. In this study, building on our observation that *shs1-ΔCTD* cells are delayed in re-entering division, we examined the ER stress-induced behavior of upstream Cdc42 regulators and spindle positioning in both WT and *shs1-ΔCTD* backgrounds.

## RESULTS

### ER stress-induced localization of Bud proteins

In our previous investigations, in addition to the mis-localization of the septin ring and the small GTPase Cdc42 to the bud scar, we showed that ER stress also causes mis-localization of Cdc24—the guanine nucleotide exchange factor (GEF) for Cdc42—and the Bud1/Rsr1 GTPase to the bud scar^8^. These findings revealed that the Cdc42 polarity complex forms at the bud scar under ER stress, even though Cdc42 is inactivated rapidly. To further examine the behavior of this complex, we analyzed the localization of Bud1/Rsr1 regulatory components, including Bud2, a GTPase-activating protein (GAP), and Bud5, a guanine nucleotide exchange factor (GEF)^8^ (**Figure 1A**). As in our earlier work, we classified cells for ER inheritance by cell-cycle stage based on the mother–daughter bud index: **G1 phase**: unbudded, **S phase**: bud index < 0.33, **G2 phase**: bud index 0.33–0.67, **M phase**: bud index > 0.67¹.

**Figure 1:**
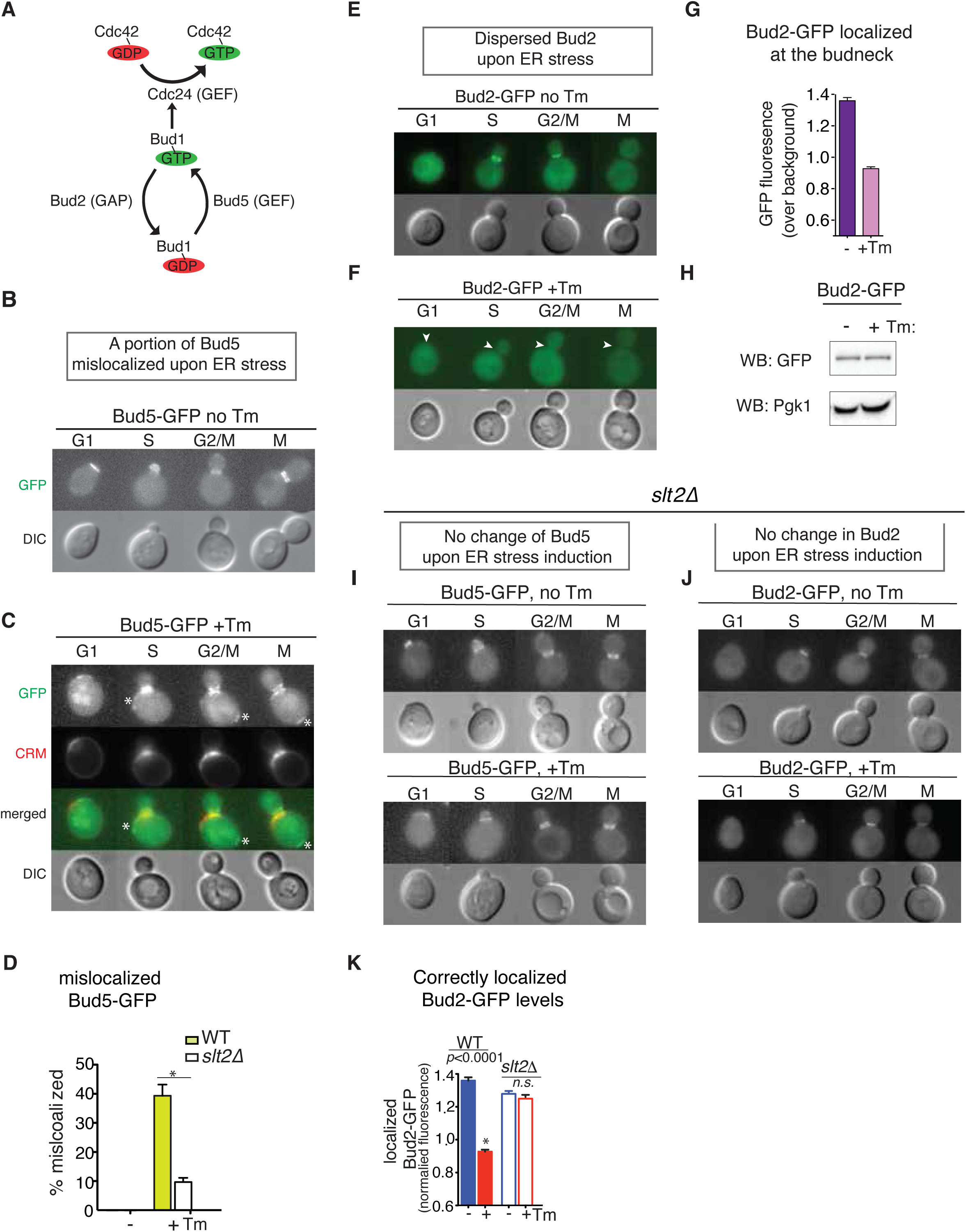
Bud2, but not Bud5, localization is altered upon ER stress in an ERSU-dependent manner. (A) Schematic of the Cdc42-activating Bud1/Rsr1 GTPase module. (B) Bud5-GFP localization in unstressed (non-Tm-treated) cells. (C) Bud5-GFP localization in Tm-treated cells. * indicates mislocalized Bud5-GFP away from bud neck. (D) Quantification of mislocalized Bud5-GFP in Tm-treated WT (pale green) and *slt2Δ* (white) cells. (E) Localization of Bud2 (Bud2-GFP) in unstressed cells. (F) Localization of Bud2 (Bud2-GFP) in Tm-treated cells. White arrowheads point lack of Bud2 at bud neck (G) Quantification of bud neck localized Bud2-GFP levels in untreated (-) (dark purple) and Tm-treated (+Tm, light purple) cells. (H) Bud2-GFP protein levels remain unchanged in WT cells treated with DMSO or Tm. Total yeast protein extract from ER stressed and unstressed cells were analyzed for the Bud2-GFP protein levels by western blot analysis probing with anti-GFP antibody. Similar levels of Pkg1 protein serves as a loading control. (I) Bud5-GFP localization in *slt2Δ* cells, with and without Tm treatment. (J) Bud2-GFP localization in *slt2Δ* cells, with and without Tm treatment. (K) Quantification of Bud2-GFP levels in WT and *slt2Δ* cells.

Under unstressed conditions, Bud5 localized to the site of polarized growth (**Figure 1B**). Upon ER stress induction with tunicamycin (Tm)—an inhibitor of N-glycosylation that triggers ER stress— Bud5 re-localized to the bud scar or cytokinetic remnants (CRMs) in ∼40% of stressed cells (**Figures 1C–D**). Similarly, under normal growth, Bud2 was localized to the site of polarized growth in G1 phase and concentrated at the bud neck during S and G2/M phases as reported previously (**Figure 1E**). Following ER stress, Bud2 lost its defined localization, becoming dispersed throughout the cell (**Figure 1F**), despite total Bud2–GFP levels remaining unchanged between unstressed and stressed cells (**Figures 1G–H**). This dispersion indicates that Bud2 is no longer concentrated at the bud neck, at CRMs, or at any other specific subcellular site. Importantly, both Bud5 mislocalization to the bud scar and Bud2 dispersal under ER stress require SLT2, a central component of the ER surveillance (ERSU) pathway. In *slt2Δ* cells, Bud5 and Bud2 maintained their normal localization at the bud neck even under ER stress, similar to their distribution in unstressed cells (**Figures 1D, 1I–K**), suggesting the idea that ER stress–induced mis-localization of Bud2 and Bud5 occurs as ERSU-dependent events.

### Impact of ER stress on the ER diffusion barrier

Previous studies reported that the septin subunit Shs1 contributes to establishing the ER diffusion barrier, and that a C-terminal truncation of Shs1 (*shs1-ΔCTD*) weakens this barrier*^13, 14^*. Furthermore, we previously showed that in ER-stressed *shs1-ΔCTD* cells, active Cdc42 is absent, while Cdc24 is mislocalized to the CRMs, albeit at reduced levels^8^. To further investigate, we examined Bud5 localization in both unstressed and ER-stressed *shs1-ΔCTD* cells. Bud5 remained at the site of polarized growth throughout the cell cycle under both conditions (**Figures *S1A–B***). In contrast, Bud2 was dispersed in *shs1-ΔCTD* cells regardless of ER stress status (**Figures *S1C–E***), suggesting that its localization depends on an intact ER diffusion barrier. Given that Bud2 has been implicated in the spindle position checkpoint (SPoC)^15^—which ensures proper spindle alignment before mitotic exit—disruption of the ER diffusion barrier may contribute to defective spindle positioning.

### ER stress accelerates spindle pole body replication and positioning

The influence of ER stress on Bud2 localization and its implications for spindle positioning prompted us to investigate if ER stress affects spindle pole body (SPB) dynamics, focusing on both the timing of SPB duplication and its spatial orientation^16-18^. To this end, we employed Spc110 tagged with GFP (Spc110-GFP), an established marker of the SPB’s inner plaque, as a biosensor to monitor these processes in live cells.

In unstressed conditions, a single Spc110-GFP in unbudded mother cells duplicated during S phase (**Figure 2A-D, *S2A***). Following duplication, the new SPB was inserted into the nuclear envelope (NE) in close proximity to the original SPB (**Figure 2A-D**, S phase)^19^. As the cell cycle progressed, one of the Spc110-GFP foci migrated to the opposite side of the NE, forming a short bipolar spindle characteristic of the G2 phase (**Figure 2A-D**, G2/M phase). This was followed by nuclear migration into the daughter cell and subsequent nuclear division (**Figure 2B**, M phase). Notably, under these normal growth conditions, unbudded cells never displayed two distinct Spc110-GFP foci (**Figure 2B-C**). Upon induction of ER stress with tunicamycin (Tm), however, we observed a marked shift in SPB dynamics. SPB duplication occurred prematurely, with two Spc110-GFP foci frequently present in unbudded G1-phase cells (**Figures 2B-D**, G1, +Tm). Furthermore, one Spc110-GFP was transferred to the daughter cell in ER stressed G2/M cells. A similar phenomenon was seen when cells were treated with dithiothreitol (DTT), another potent ER stress inducer, resulting in approximately half of unbudded cells exhibiting duplicated SPBs (**Figure 2C**, +DTT). These findings revealed that ER stress accelerates the timing of SPB duplication, uncoupling it from its usual cell cycle constraints (**Figure 2C-D**). Beyond altered duplication kinetics, ER stress also disrupted the spatial positioning of SPBs. In stressed cells, one of the duplicated Spc110-GFP foci entered the daughter cell even when the bud was still small, corresponding to the S phase (**Figure 2B**, +Tm, * marks an extra Spc110-GFP foci). By contrast, in unstressed cells, the second SPB typically appeared in the daughter cell during the M phase, following a well-defined temporal sequence (**Figure 2B and 2D, and *S2A***), consistent with previous report (reviewed in ^20^). The premature entry of the SPB into the daughter cell under ER stress suggests a breakdown in the mechanisms that normally coordinate spindle orientation with cell cycle progression. To further validate these observations, we utilized an additional SPB biosensor, Spc42 fused to RFP (Spc42-RFP) (**Figure 2E**). The slower folding kinetics of RFP allowed us to distinguish newly synthesized SPBs (which appeared smaller and dimmer) from pre-existing ones^21^. Under normal conditions, the newly generated SPB (with dimmer RFP) was retained within the mother cell, as previously reported (**Figure 2E**, +DMSO, M phase)^22-24^. However, ER stress led to an increased frequency of a dimmer Spc42-RFP signal in daughter cells (**Figure 2E**, +Tm, M phase), indicating that the asymmetric retention of the new SPB in the mother was compromised.

**Figure 2:**
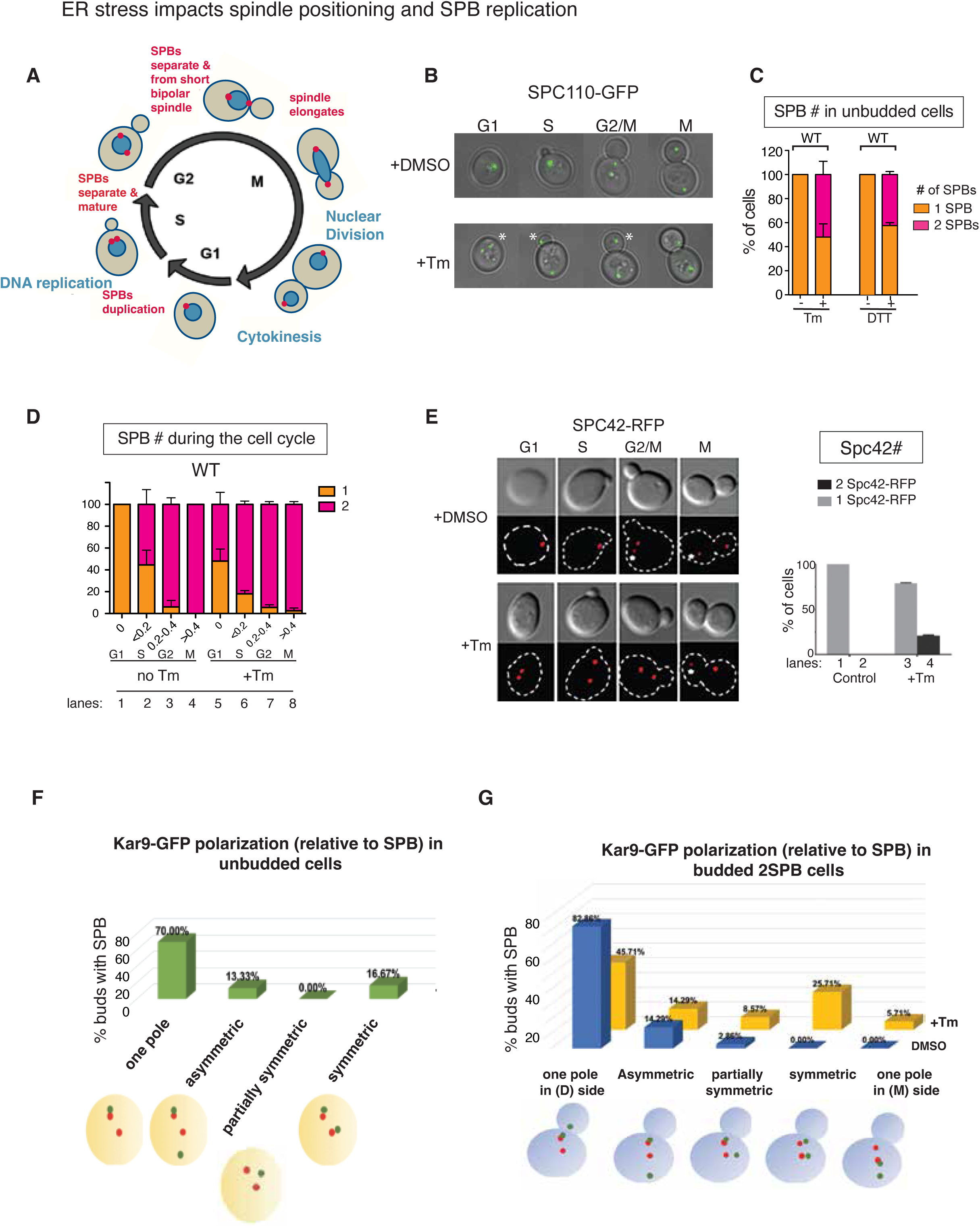
ER stress accelerates the kinetics of SPB duplication and spindle positioning. (A) Graphic representation of the cell cycle stages of yeast, *S. cerevisiae*. In the G1 phase, a single SPB (red foci) is located on the nuclear membrane (dark green). As the daughter cell emerges and enters the S phase, SPB duplication occurs, with the newly generated SPB inserted into the NE adjacent to the pre-existing SPB. During G2, one SPB migrates to the opposite side of the NE, establishing a bipolar spindle. As mitosis progresses, spindles elongate, enabling nuclear entry into the daughter cell and nuclear division. Finally, nuclear division separates the two SPBs into two nuclei before cytokinesis occurs, producing two daughter cells. **(B)** SPB duplication, positioning, and separation were monitored using an SPB biosensor, Spc110-GFP, in unstressed (DMSO-treated) and Tm-treated cells (1 μg/ml Tm for 2.5 hr). Cell cycle stages (G1, S, G2/M, and M) were determined by bud index (ratio of daughter cell diameter to that of the mother cell), as described^8^. G1: unbudded cells (bud index = 0), S: bud index <0.2, G2/M: bud index 0.2–0.4, and M: bud index >0.4. **(C)** Quantification of WT unbudded cells with either one SPB (orange) or two SPBs (magenta), either untreated (lanes 1 and 3) or treated with Tm (lane 2) or DTT (lane 4). **(D)** Quantification of WT cells with either one SPB (orange) or two SPBs (magenta) at different cell cycle stages, either untreated (lanes 1-4) or treated with Tm (lane 5-8). **(E)** *Spc42*, another SPB subunit fused to RFP, exhibited accelerated duplication in response to ER stress (+Tm). White circle points SPB with diminished level of RFP, suggesting the newly generated SPB based on slow folding of RFP. Quantification showed a significant increase in unbudded WT cells with duplicated SPBs (dark gray) only in Tm-treated cells. White closed circle indicates a SPB with diminished RFP level, suggesting the newly generated SPB. **(F)** Quantification of Kar9-GFP (green circles) localization and positioning relative to replicated SPBs (red circles) in unbudded cells upon ER stress induction. **(G)** Quantification of Kar9-GFP (green circles) localization and positioning relative to replicated SPBs (red circles) in budded cells under DMSO and Tm treated conditions.

Under unstressed conditions, SPB duplication and asymmetry are tightly coordinated with activation of Cdc28/Clb5 in the G2/M phase. Early in the cell cycle, the pre-existing SPB associates with Kar9 before the appearance of the new SPB^23, 25, 26^. The newly synthesized SPB emerges later, when Cdc28/Clb5 activity rises and phosphorylates the new SPB, which prevents association with Kar9. Thus, the kinetic orders of events ensure that only the pre-existing SPB is initially exposed to Kar9, thereby establishing SPB asymmetry and directing proper spindle orientation. ER stress induction, by accelerating SPB duplication, causes both the pre-existing and new SPBs to be present early in the cell cycle stages and simultaneously exposed to Kar9. This premature exposure undermines the establishment of SPB asymmetry, potentially compromising the fidelity of spindle positioning and subsequent chromosome segregation. Collectively, these results highlight a critical link between ER homeostasis and the spatial-temporal regulation of the mitotic spindle apparatus, with ER stress disrupting both the timing and orientation of SPB dynamics (**Figure 2F-G**).

### ER stress also impacts spindle orientation

The duplication of spindle pole bodies (SPBs) is crucial for the attachment and orientation of mitotic spindles during cell division. To determine how ER stress-induced changes in SPB behavior—particularly the loss of SPB asymmetry—affect microtubule organization, we used Tub1-GFP to visualize spindle microtubules (**Figure 3A–B**). We tracked the angle (θ) between the spindle axis (defined by the two SPBs) and the mother–daughter cell axis throughout the cell cycle (**Figure 3A**). In wild-type cells under normal growth, θ initially measured 50°–90° in early stages (bud index <0.2) and gradually decreased as cells progressed, reaching 0°–20° in late stages (bud index >0.4), reflecting precise alignment with the bud axis^27^ (**Figure 3A–B**). Under ER stress, however, θ in small-budded cells began at a much lower value—similar to that of medium-budded unstressed cells—and changed little thereafter (**Figure 3B**, WT), indicating a substantial loss of cell cycle–coupled spindle rotation. Strikingly, shs1-ΔCTE mutant cells displayed a reduced θ from early stages onward, regardless of stress, consistent with a persistent orientation defect (**Figure 3B**).

**Figure 3:**
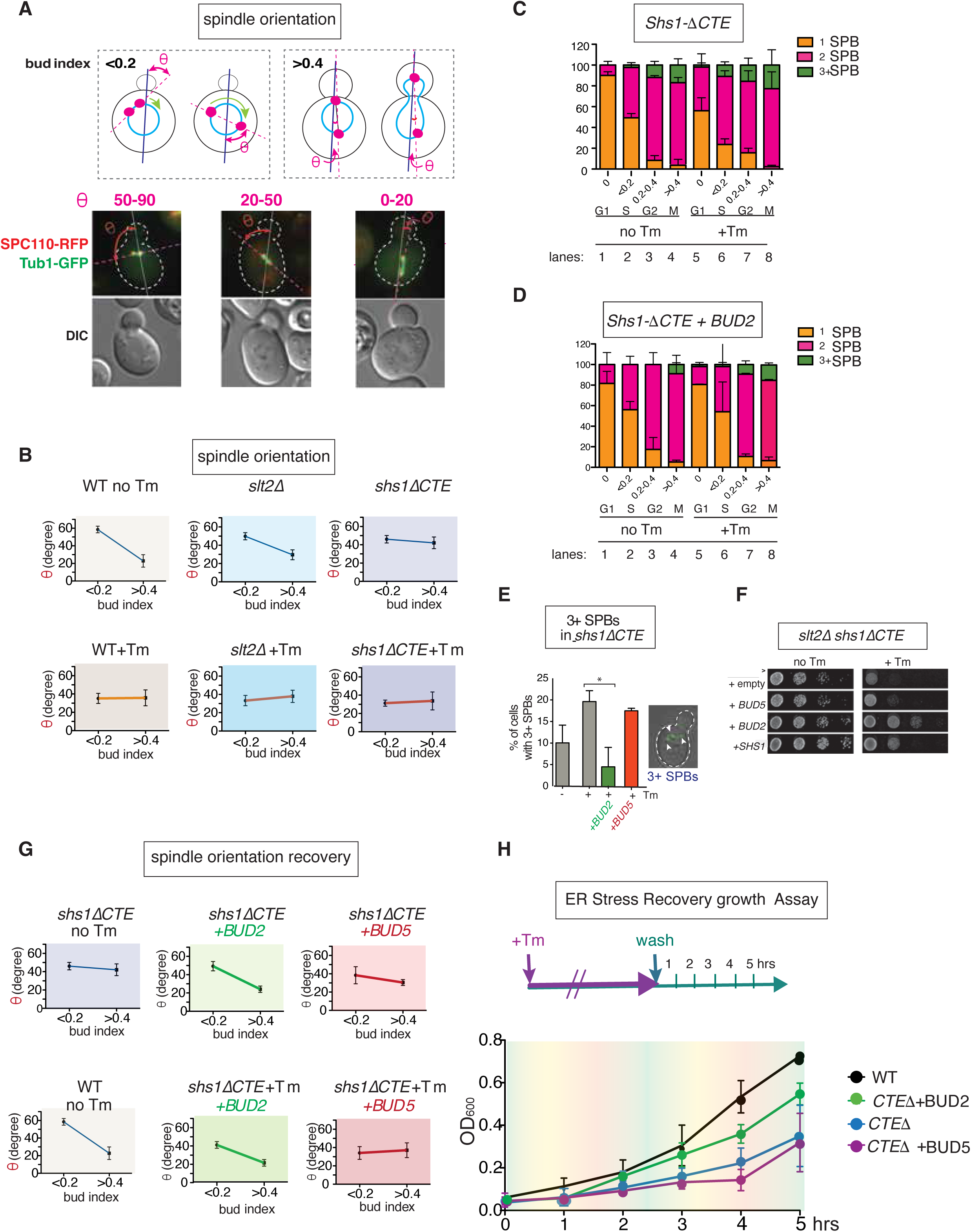
ER stress accelerates spindle pole body duplication and alters spindle assembly kinetics. (A) Spindle positioning, defined by two SPBs (magenta circles), was analyzed in small-budded cells (bud index <0.2) and large-budded cells (bud index >0.4) relative to the mother-daughter cell axis. The angle (θ) between these axes is shown. Images of cells expressing SPC110-RFP and Tubulin-GFP (Tub1-GFP) illustrate spindle positioning. **(B)** Average values of θ at early (<0.2) and late (>0.4) cell cycle stages in WT, *slt2Δ* and *shs1-ΔCTE* cells, shown for unstressed (no Tm) and ER-stressed (+Tm) conditions. The bud index was used to define cell cycle stages: <0.2 represents the early stage (G1/S phase) with small buds, while >0.4 corresponds to later stages (G2/M phase) with large buds. **(C)** SPB duplication, positioning, and separation were monitored using an SPB biosensor, Spc110-GFP, in unstressed (DMSO-treated) and Tm-treated *shs1-*Δ*1CTE* cells (1 μg/ml Tm for 2.5 hr). Cell cycle stages (G1, S, G2/M, and M) were determined by bud index (ratio of daughter cell diameter to that of the mother cell), as described in Chao et al., 2019^8^. G1: unbudded cells (bud index = 0), S: bud index <0.2, G2: bud index 0.2–0.4, and M: bud index >0.4. **(D)** Quantification of SPBs in *shs1-ΔCTE* cells overexpressing *BUD2* at different cell cycle stages, with or without Tm treatment under untreated (no Tm, lanes 1–4) and Tm-treated (lanes 5–8) conditions. **(E)** Numbers of cells with 3 SPBs in ER stressed *shs1-Δ1CTE* was diminished by overexpression of BUD2, but by that of BUD5. An image of a *shs1-ΔCTE* cell with three SPBs is shown in a small inset. **(F)** Growth of ER stressed *slt2Δ1shs1-Δ1CTE* cells was rescued by BUD2, but not BUD5, overexpression, closer to SPB duplication in unstressed *shs1-Δ1CTE* cells. **(G)** Average values of θ at early (<0.2) and late (>0.4) cell cycle stages in *shs1-ΔCTE* cells overexpressing *Bud2* or *Bud5*. *Bud2* overexpression restored θ in both unstressed and ER-stressed *shs1-ΔCTE* cells similar to those of unstressed WT cells, while *Bud5* overexpression did not. **(H)** Schematic of the ER stress recovery and re-entry assay into the cell cycle for ER-stressed *shs1-ΔCTE* or *shs1-ΔCTE* cells overexpressing *Bud2* or *Bud5*. To facilitate ER stress recovery, cells were washed and grown in media without Tm. The impact of *Bud2* and *Bud5* was examined by transforming *shs1-ΔCTE* cells with plasmids expressing *Bud2* or *Bud5* under the strong *TEF* promoter. The growth of *shs1-ΔCTE* cells re-entering the cell cycle was monitored over time.

Because Bud2 was diffused in both unstressed and ER stressed *shs1-ΔCTE* cells (**Figure *S1A–C***), we tested whether its dispersal contributed to abnormal SPB duplication and positioning. Bud2’s failure to concentrate at the bud neck, a site critical for the spindle assembly checkpoint, correlated with duplication defects: some unbudded shs1ΔCTE cells already contained two SPBs on the nuclear envelope (NE) (compare **Figure 3C vs 2D**), and others developed three SPBs even without ER stress (**Figure 3C**, lanes 3-4). ER stress exacerbated these phenotypes, with more S phase *shs1ΔCTE* cells displaying three SPBs (**Figure 3C**, lanes 6–8), suggesting severely disrupted duplication kinetics and fidelity.

Bud2 localization was specifically altered by ER stress in WT cells (**Figure 1F**) and *shs1ΔCTE* cells, while Bud5—a GEF for Bud1/Rsr1 and another Cdc42 effector—remained largely unaffected by ER stress (**Figure *S1D–E***). Overall Bud2 protein levels did not change (**Figure 1H**), suggesting that defects arose from Bud2 mis-localization rather than reduced abundance. Overexpressing Bud2 in shs1ΔCTE cells affected overall duplication kinetics and significantly decreased the proportion of cells with ≥3 SPBs under both stressed and unstressed conditions (**Figure 3D-E, and 3G**). We found that this intervention modestly affected overall SPB duplication kinetics in both unstressed (**Figure 3C**, lanes 1–4 vs. **Figure 3D**, lanes 1–4) and ER-stressed cells (**Figure 3C**, lanes 5–8 vs. **Figure 3D**, lanes 5–8), and notably reduced the proportion of cells with three or more SPBs, both in unstressed and ER-stressed conditions (**Figure 3C-E**). In contrast, Bud5 overexpression neither corrected duplication kinetics nor reduced supernumerary SPB formation under ER stress (**Figure 3E**). Furthermore, Bud2—but not Bud5—overexpression restored spindle orientation in shs1ΔCTE cells under both conditions to levels comparable to unstressed WT and restored cell growth of slt2Δ1shs1ΔCTE cells (**Figure 3F-G**).

Previously, we showed that shs1-ΔCTE cells mount hallmark ERSU responses, blocking ER inheritance and cytokinesis under ER stress, but failed to re-enter the cell cycle upon restoration of ER homeostasis^8^. Using our ER stress recovery assay (**Figure 3H**), we found that Bud2 overexpression markedly accelerated the cell cycle re-entry kinetics in *shs1-ΔCTE* cells, to levels closer to WT, whereas Bud5 overexpression had little effect. These results suggests that Bud2 plays as a critical regulator linking ER homeostasis to spindle dynamics, nuclear events, and promoting timely cell cycle progression upon restoration of ER function, likely by mediating the ER stress–induced re-localization of Shs1, Cdc42, and upstream effectors to bud scars.

### ERSU pathway regulates ER stress-induced spindle phenotypes

The ERSU pathway coordinates key responses to ER stress, including blocking ER inheritance, relocalizing the Shs1 septin ring subunits to the bud scar, and arresting cytokinesis. To determine whether this pathway also regulates spindle pole body (SPB) dynamics—specifically, duplication timing and translocation into the daughter cell during ER stress—we examined the contributions of ERSU genes such as **RTN1** and **YOP1**. We reported previously that the ER membrane domains of these proteins contain motifs that bind phytosphingosine (PHS), an early intermediate in sphingolipid biosynthesis, which is required for activating both the ER inheritance block and septin ring mislocalization during ER stress^5, 6^. Mutations such as **Rtn1-L56A**, which disrupt PHS binding, abolish the ER inheritance block without altering overall ER morphology. Notably, ER stress–induced acceleration of SPB duplication was reduced in unbudded *rtn1Δrtn2Δyop1Δ* cells expressing Rtn1-L56A (**Figure 4A**), supporting the idea that these spindle duplication phenotypes are, at least in part, mediated by the ERSU pathway. By contrast, *rtn1Δrtn2Δyop1Δ* cells expressing **Rtn1-Y60A**, which also impairs PHS binding, displayed SPB duplication comparable to that of cells expressing WT Rtn1, suggesting that additional components beyond PHS–Rtn1 binding contribute to spindle regulation.

**Figure 4:**
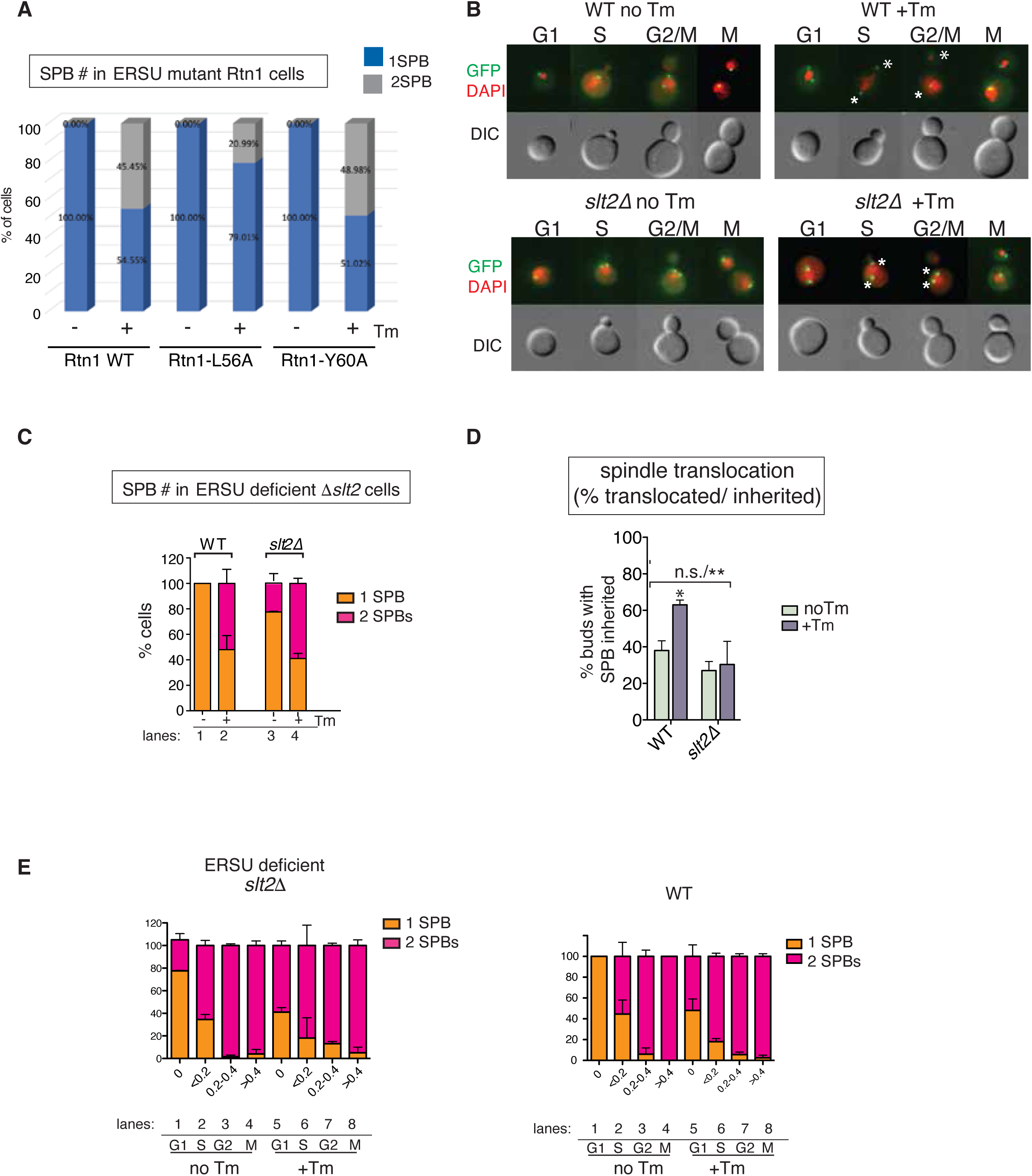
ER stress-induced SPB duplication and positioning phenotypes are part of the ERSU pathway. (A) Number of unbudded cells with one or two SPBs in unstressed and ER-stressed *rtn1Δrtn2Δyop1Δ* cells expressing Rtn1-WT or ERSU deficient alleles of Rtn1, Rtn1-L56A or Rtn1-Y60A mutant cells with (+) or without (-) Tm. **(B)** SPBs duplication and translocation to the daughter cell in unstressed (no Tm) and ER-stressed (+Tm) unbudded WT and ERSU deficient *slt2Δ* cells. SPB was monitored by SPC110-GFP and nuclear DNA was stained by DAPI. White stars in ER stressed S and G2/M WT cell show that one of the two SPBs was translocated into the daughter cell, while both remained in S and G2/M *slt2Δ1* mother cells. **(C)** Number of SPBs in unbudded unstressed (no Tm) and ER-stressed (+Tm) WT and *slt2Δ* cells. (F) The percentage of WT, or *slt2Δ* cells with translocated SPBs in the bud was measured under-Tm and +Tm conditions. (G) SPB duplication in unstressed and ER-stressed *slt2Δ* cells throughout cell cycle. For comparisons, the SPB duplication result from WT cells is taken from Figure 2D.

We also investigated the role of SLT2, another ERSU component, in SPB duplication and positioning—particularly the translocation of SPBs into daughter cells (**Figure 4B-D**). In the absence of ER stress, *slt2Δ* cells showed small increase in unbudded and S-phase with two SPBs (**Figure 4B and C**), suggesting that accelerated SPB duplication could occur independently of SLT2. However, in *slt2Δ* cells, the frequency of SPB translocation into the daughter cell was not significantly altered by ER stress. In wild-type cells, ER stress markedly accelerated SPB entry into the daughter cell during S phase, but this effect was absent in *slt2Δ* cells, where translocation kinetics in both unstressed and stressed conditions remained similar. These findings identify SLT2 as a regulator of SPB translocation into the daughter cell under ER stress (**Figure 4B and 4E**), while implicating the broader ERSU pathway in both SPB positioning and, to a lesser extent, ER stress–induced acceleration of SPB duplication.

### ER diffusion barrier defects impair nuclear entry into the daughter cell and nuclear division

The effects of ER stress on SPB translocation into the daughter cell suggested a potential influence on nuclear positioning and division. Upon association with Kar9, SPBs migrate along microtubules toward the daughter cell, coordinating the spatial dynamics of mitosis. To examine how ER stress impacts perinuclear ER (pnER) inheritance and nuclear division, we monitored the distribution of the established pnER marker Pho88-GFP (**Figure 5A**). This approach allowed us to classify cells based on pnER localization—whether retained in the mother or daughter—and to distinguish between cells with or without completed nuclear division. We also identified cells in which pnER was positioned between the mother and daughter (“dividing”) and those in which the divided nuclei were retained entirely within the mother cell (“error”) (**Figure 5A**).

**Figure 5:**
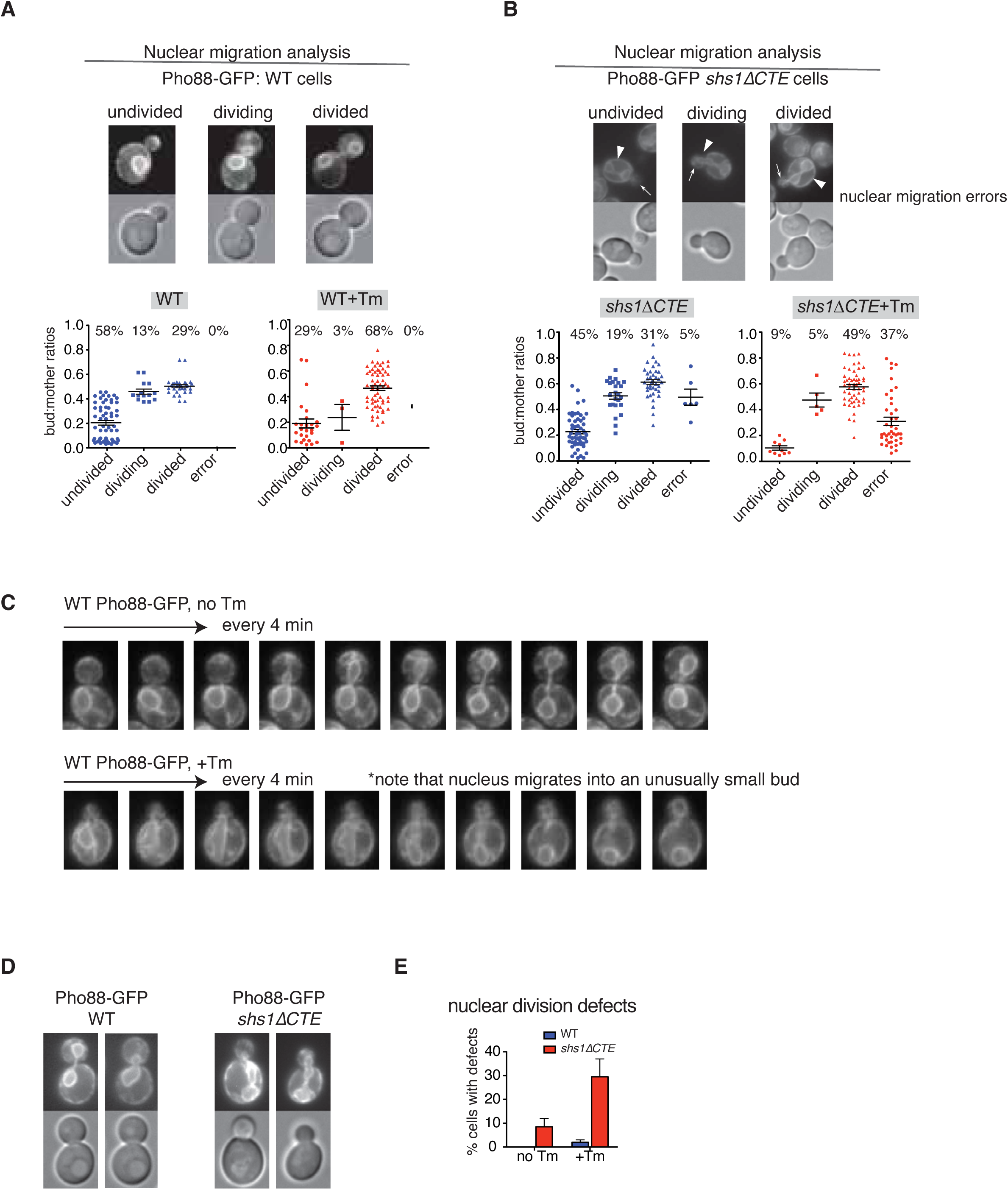
ER stress compromises nuclear transfer into the bud and nuclear division. (A) Nuclear migration and division in WT cells carrying *Pho88-GFP* with (+Tm) or without (-Tm) ER stress induction. Examples of undivided, dividing, and divided cell images are shown. Quantitation of bud:mother ratios for WT was shown for Tm treated or untreated cells. **(B)** Nuclear migration and division in *shs1-ΔCTE* cells carrying *Pho88-GFP* with (+Tm) or without (-Tm) ER stress induction. Examples of undivided, dividing, and divided cell images are shown. Quantitation of bud:mother ratios for *shs1-ΔCTE* was shown for Tm treated or untreated cells. **(C)** Time-lapse (ever 4 minutes) images of WT cells carrying Pho88-GFP with or without Tm treatment, showing nuclear transfer into the daughter cell in ER stressed WT cell occurs when the daughter cell is small. **(D)** Representative images of unstressed and stressed WT and *shs1Δ1-CTE* cells carrying Pho88-GFP **(G)** Quantification of nuclear division in WT and *shs1-ΔCTE* cells treated with or without Tm.

In asynchronous wild-type populations, most cells (58%) displayed undivided pnER confined to the mother cell (**Figure 5A**, WT). Following ER stress, this distribution shifted markedly, with 68% of cells exhibiting “divided” pnER, consistent with accelerated SPB-related processes such as duplication, positioning, and transfer. In contrast, unstressed *shs1-ΔCTE* mutants showed slightly elevated baseline levels of both “divided” (31%) and “dividing” (19%) pnER compared with unstressed wild type (29% and 13%, respectively), even before stress induction (**Figure 5B**). Notably, a fraction of unstressed *shs1-ΔCTE* cells already exhibited erroneous nuclear division, retaining divided nuclei in the mother cytoplasm—a defect that increased to 37% under ER stress (**Figure 5B**). These findings indicate that ER stress compromises the pnER diffusion barrier, leading to misregulation of pnER, nuclear. However, even in *shs1-ΔCTE* cells lacking a robust ER diffusion barrier, the ER inheritance block triggered by ER stress remained intact, as confirmed by additional analyses (**Figure *S3A***). Time-lapse imaging revealed that, under normal conditions, wild-type cells completed nuclear division with accurate segregation between mother and daughter cells (**Figure 5C**). Under ER stress, however, nuclear entry into the daughter cell occurred prematurely, sometimes when the bud was still small, and nuclear division occasionally took place entirely within the mother cytoplasm. Quantitative analysis confirmed that ER stress increased the frequency of uneven nuclear division defects in wild-type cells (**Figure 5D-E**, WT + Tm), with these defects becoming even more pronounced in *shs1-ΔCTE* mutants (**Figure 5D-E**, *shs1-ΔCTE*). Together, these results suggest that the ER diffusion barrier is essential for proper pnER inheritance.

### Altered nuclear division causes uneven chromosome division

Disruptions in nuclear entry and division can lead to unequal distribution of nuclear DNA between mother and daughter cells. Because daughter cells generally grow more slowly than mother cells, their overall size—and consequently their nuclear size—is typically smaller, as previously documented (**Figure 6A**, no Tm, blue arrow). Upon induction of ER stress in wild-type cells, DAPI staining revealed an even greater reduction in the DNA-stained area within daughter nuclei (**Figure 6A**, +Tm, white arrow). Quantitative measurements showed that, under unstressed conditions, the ratio of DAPI fluorescence intensity between mother and daughter nuclei was close to 1.0, consistent with comparable DNA content. Given that ER stress reduced daughter cell size, we normalized nuclear DNA measurements to cell diameter, determined from DIC imaging. This normalized ratio fell significantly following tunicamycin treatment, indicating a marked decrease in DNA content in daughter cells (**Figure 6B**).

**Figure 6:**
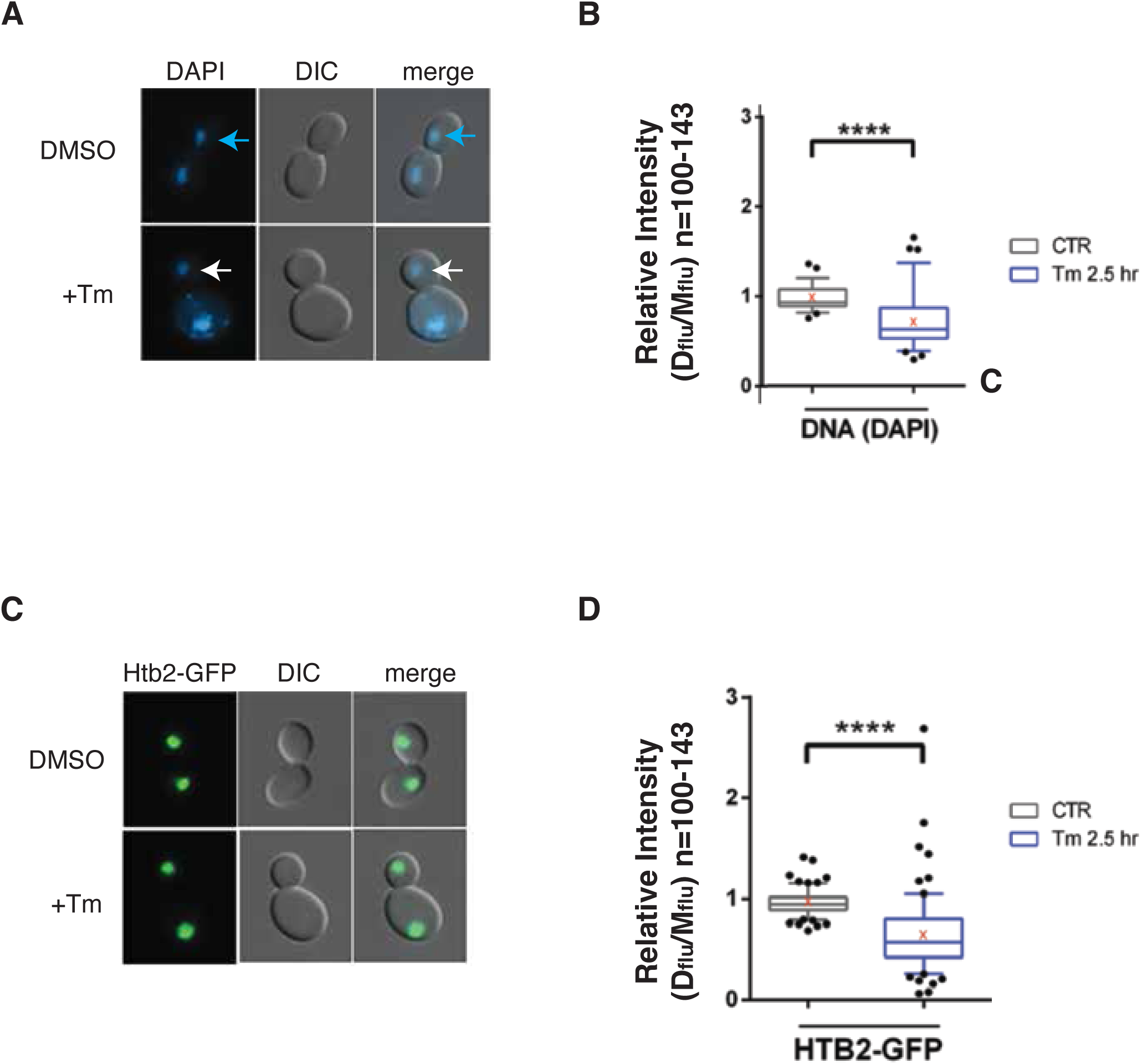
ER stress impacts nuclear DNA division in daughter cells. (A) Images of nuclear DNA stained with DAPI in ER-stressed (+Tm) and unstressed (-Tm) WT cells were shown. **(B)** Quantification of DAPI in cells shown in (A) with or without Tm treatment. Mother and daughter cell sizes were adjusted (DIC). P < 0.01. **(C)** Images of Histone H2B, monitored by Htb2-GFP in mother and daughter ER-stressed (+Tm) and unstressed (-Tm) cells, were shown. **(D)** Quantitation of Htb2-GFP with or without Tm with corresponding quantification. P < 0.01.

To validate these findings, we compared DAPI-based DNA measurements (**Figure 6A**) with histone H2B-GFP (Htb2-GFP) signals (**Figure 6C**) in M-phase cells after nuclear division. Normalization of Htb2-GFP fluorescence to daughter cell size confirmed that daughter cells exposed to ER stress retained less DNA than those in unstressed conditions (**Figure 6B & 6D**). Together, these results demonstrate that ER stress compromises the fidelity of DNA segregation, leading to an unequal distribution of nuclear DNA between mother and daughter cells.

## DISCUSSION

The successful formation of progeny cells hinges on the precise coordination of nuclear segregation and organelle distribution throughout the cell cycle, spanning from initiation to cytokinesis^28-32^. Our studies demonstrate that this coordination falters when the functional homeostasis of a single organelle, such as the endoplasmic reticulum (ER), is compromised. Specifically, impairment of ER homeostasis blocks ER inheritance and disrupts nuclear cell cycle events by causing misorientation of mitotic spindles within the mother cell. Moreover, ER stressed cells transfer the spindle and nucleus to daughter cells in an uncontrolled fashion. Consequently, daughter nuclei in ER-stressed cells often appear smaller with reduced DAPI staining, resulting in uneven DNA division and diminished DNA content. These findings reveal that disrupted ER functional homeostasis perturbs nuclear cell cycle regulation and ultimately blocks cytokinesis, partly due to the septin ring subunit, Shs1, translocating to bud scars. This positions cytokinesis as a crucial organizational checkpoint, ensuring coordination between organelle function and nuclear events^33-35^.

Our work uncovers that molecular events triggered by ER homeostasis are closely linked to nuclear cell cycle processes in yeast, particularly those governing spindle positioning and entry into the daughter cell. Similarly, the link of organelle inheritance with cell cycle progression has been described for vacuole/lysosome^32,36^. Furthermore, previous reports indicate that genetic disruptions impairing peroxisome inheritance also cause spindle misalignment^37^, paralleling our observation that blocked ER inheritance due to ER stress leads to spindle orientation errors in yeast. Although our findings originate from yeast studies, analogous regulatory mechanisms likely exist in mammalian cells, including stem cells that divide asymmetrically. For example, aged mitochondria are selectively partitioned into daughter cells during mitosis in mammalian stem cells^38^, and peroxisome inheritance in skin stem cells closely correlates with spindle orientation^37^. An anticipated player emerged as a Lynchpin between the ER and mitotic DNA turned out to be Bud2, one of the polarity establishment components after ER stress-induced inactivation. Notably, our recent data revealed that ER stress in mammalian cells affects metaphase by altering positioning of chromatin condensation, spindle positioning, and orientation with respect to the metaphase ER (Shiozaki et al., in preparation), suggesting that the molecular mechanism that separates functional ER and mitotic chromatin are intimately orchestrated.

### Septin Dynamics in the ERSU Cell Cycle Checkpoint

Our results indicate that the ERSU pathway monitors both ER function and spindle checkpoints, with the septin subunit Shs1 playing a lynchpin. Initially identified as essential for cell division, septins now are recognized for broader functions including establishing cell polarity ^39-41^ ^42-46^, scaffolding the cytoskeleton, forming diffusion barriers in the plasma membrane and ER, and defining daughter cell identity during bud emergence^47^. These roles generally require septins to serve as stable landmarks through much of the cell cycle. However, under ER stress, we observe an unprecedented relocation of the Shs1 septin subunit from the bud neck to bud scars. This re-localization may activate the Swe1-dependent morphogenesis checkpoint^48, 49^, delaying mitosis and effectively halting cell cycle progression—likely providing the cell with extra time to alleviate ER stress.

Mislocalization of Shs1 also disrupts Bud2 positioning, leading to two major consequences. First, as Bud2 acts as the GAP for Bud1/Rsr1 (a GTPase upstream of Cdc42), its mislocalization impairs Cdc42 activation for the polarized growth, thereby blocking polarized growth. Second, because Bud2 is part of the spindle position checkpoint (SPoC)^15^, its misplacement permits ER-stressed cells to translocate misoriented spindles. Critically, spindles entering the bud frequently carry smaller nuclei, suggesting incomplete genome duplication or uneven chromosome segregation. Previously, we confirmed that DNA replication proceeds normally with sister chromatids assembled correctly when ER stress is induced^50^. Hence, our findings suggest that cells bypass SPoC under ER stress via Bud2 mislocalization, enabling nuclear transfer into stressed daughter cells. Because ER-stressed cells fail to complete division and resume growth only after ER function restoration, this may mark daughter cells as repositories for damaged nuclear material. Supporting this, we reported that initial daughter cells often remain attached to the mother after ER stress recovery^1^. Furthermore, recovery of the ER functional homeostasis leads to a new bud emerging from the mother cell, ultimately undergoing cell division. Future work must elucidate the molecular mechanisms by which restoring ER homeostasis facilitates cell cycle re-entry.

The importance of disassembling the Bud1/Rsr1 complex and inactivating polarized Cdc42 during ER stress is highlighted by the phenotypes of slt2Δ and slt2Δ shs1ΔCTE mutants^8^. In ER-stressed slt2Δ cells, Bud2 and/or Bud5, which act upstream of Bud1/Rsr1 and CDC24, fail to respond to ER stress, causing poor survival. By contrast, in slt2Δshs1ΔCTE mutants, these proteins mis-localize even without stress, impairing growth and increasing stress sensitivity. Normally, Cdc42 regulation maintains singular budding sites; loss of this control leads to multiple buds, as seen in these mutants^8^. Importantly, Bud2 overexpression or Cdc42 inhibition partially rescues these defects, underscoring the integral role of polarity regulation in the ER stress response.

Is there a safeguard preventing stressed daughter cells from sustaining growth or recovering polarity? We propose two possibilities. First, the ERSU pathway blocks cortical ER (cER) inheritance into daughter cells during ER stress; since cER inheritance is a stepwise process, stressed daughters may reach a “point of no return” after which cER cannot return back to the mother cell. Second, polarity establishment relies on competition among Cdc42 activation foci, with positive feedback reinforcing dominant sites^51^. During ER stress, core Cdc42 components—including septins, Cdc24, and Bem1—cluster at bud scars^8^. Re-establishing polarity requires relocation outside this inhibitory zone, making new buds more likely to form on mother than on stressed daughter cells. Indeed, cell observations during ER stress recovery show new buds frequently emerge adjacent to CRMs, mimicking typical axial budding patterns. Mutants defective in septin ring translocation to CRMs recover poorly and exhibit daughter cell positioning errors, supporting this model.

Our findings establish that maintaining ER functional integrity is vital for coupling organelle inheritance with nuclear genome positioning and stability. Disruption of ER homeostasis uncouples spindle orientation from organelle surveillance, causing asymmetric nuclear division and chromosome mis-segregation. While cytokinesis arrest functions as a checkpoint to limit propagation of these errors during ER stress, stressed cells can override spatial constraints via septin-mediated polarity rewiring, transferring damaged nuclear material into daughters. This coordination between organelle status and mitotic fidelity parallels conserved mechanisms in mammalian stem cells, where asymmetric organelle inheritance influences cell fate decisions^52^.

Our work broadens the role of the ERSU pathway beyond stress mitigation to directly coordinating organelle inheritance with genome dynamics. Mechanistically, the ER diffusion barrier, regulated by SLT2 and septins, organizes polarity GTPase networks to ensure proper spindle alignment during division. These insights prompt intriguing questions about whether mammalian ER stress similarly repurposes septin structures to bypass mitotic checkpoints— potentially illuminating mechanisms behind stress-induced aneuploidy in cancer cells. By defining ER inheritance as a gatekeeper of nuclear genome stability, this research reframes organelle surveillance as a cornerstone of cell cycle regulation and opens new investigative paths into how cellular stress impacts genomic integrity.

## Acknowledgements

We thank for providing the yeast strains and Drs. Douglass Forbes, Randy Hampton, and Jim Wilhelm for discussions throughout the course of this study. This work was supported by NIH R01GM087415, the California Research Cancer Coordination Fund, Paul Allen Distinguished Investigator award and UCSD Health Science Research Grant.

## Materials and Methods

### Yeast Strains and Plasmid Construction

Epitope tagging of endogenous proteins was performed via homologous recombination using PCR-amplified fragments derived from appropriate plasmid templates. Tagging was carried out at the C-terminus of the target genes at their native loci in the BY4741 haploid yeast strain. Strains containing double epitope tags (e.g., GFP/RFP) or double gene deletions were constructed using standard yeast genetic techniques, including mating, sporulation, and tetrad dissection.

Epitope-tagging of endogenous proteins was performed as described previously^8^. Briefly, by homologous recombination of PCR-generated fragments from templates in haploids at the C-terminus of the specified genes at their endogenous locus in either BY7043^53^ (Tong and Boone, 2006) or BY4741. The templates include pKT128 (GFP::SpHIS5)^54^(Sheff and Thorn, 2004), pFA6A-pmRFP-KanMX6, pFA6A-13Myc-KanMX6^55^ (B€ahler et al., 1998), pHVF1CT and pUVF2CT^13^ (Chao et al., 2014). KanMX deletion strains were obtained from freezer stocks of the haploid yeast deletion collection (BY4741, MAT a, KanMX; Thermo Fischer). NatR deletion or truncation strains were constructed in BY7092 using p4339^53^ (Tong and Boone, 2006). All deletion and truncation strains were confirmed by PCR. For the expression of BUD2, BUD5 and SHS1, full-length ORFs were cloned from BY4741 genomic DNA and inserted into p416-TEF at XbaI/SalI, as reported previously^8^.

### Yeast Cell Growth

Unless otherwise indicated, all yeast strains used in this study is of the BY4741 background. See Table S1 for a full list of strains. All yeast was grown at 30 °C in synthetic defined medium containing yeast nitrogenous base, 2% dextrose and the appropriate amino acids for the genotype unless otherwise stated.

### Yeast Spot Assays

10-fold serial dilutions of log phase cells were spotted using a pin-frogger onto agar plates containing synthetic complete (SC) media with 2% glucose and grown for 48hr at 30 °C.

### Light Microscopy

Live yeast cells in logarithmic growth phase were imaged using a Carl Zeiss Axiovert 200M microscope equipped with a 100×/1.3 NA objective lens. Colocalization assays were performed using the following tag combinations: Spc42-RFP with Kar9-GFP; Spc110-RFP with Tub1-GFP; and Spc110-RFP with Pho88-GFP. Spindle orientation was analyzed using Spc110-RFP and Tub1-GFP. All images were acquired and processed as described previously.

### Quantification of Microscopy Images

All image quantification was conducted using ImageJ software (National Institutes of Health). For each experimental condition, a minimum of 80 cells was analyzed. Cell cycle stages were classified based on bud size:

- G1 phase: no visible bud
- S phase: bud area < 1/3 of the mother cell
- G2 phase: bud area between 1/3 and 2/3 of the mother cell
- M phase: bud area > 2/3 of the mother cell

Bud index (bud-to-mother size ratio) was calculated by tracing the perimeters of the bud and mother cell in transmission images and measuring their respective areas. Relative DAPI intensity was computed by dividing the total nuclear fluorescence signal by the nuclear area in individual daughter and mother cells. Similarly, the relative intensities of HTB2 and DAPI were determined from total nuclear fluorescence in daughter versus mother cells. In box plots, the line represents the median and the cross indicates the mean value.

### Statistical Analysis

All statistical analyses were performed using GraphPad Prism (GraphPad Software). Experiments were independently repeated three times. Quantitative data are presented as mean ± standard error of the mean (SEM). Statistical significance was assessed using Student’s *t*-test, with *p*-values reported accordingly.

## Figures & Figure Legends

**Supplemental Figure 1: Bud2, but not Bud5, localization is altered upon ER stress in *shs1-Δ1CTE* cells.**

**(A)** Bud5-GFP localization in *shs1-ΔCTE* cells without Tm. White arrows points Bud5-GFP localized at the incipient bud site (G1) and bud neck (S-M phase).

**(B)** Bud5-GFP localization in *shs1-ΔCTE* cells with Tm treatment. White arrows points Bud5-GFP localized at the incipient bud site or small bud neck (G1) and bud neck (S, G2, and M phase). Red stars show mis-localizes *shs1-ΔCTE* cells.

**(C)** Bud2-GFP localization in *shs1-ΔCTE* cells without Tm. Yellow arrowheads points loss of Bud2-GFP localized at the incipient bud site (G1) and bud neck (S-M phase).

**(D)** Bud2-GFP localization in *shs1-ΔCTE* cells with Tm treatment. Yellow arrowheads points loss of Bud2-GFP localized at the incipient bud site or small bud neck (G1) and bud neck (S, G2, and M phase).

**(E)** Quantitation of Bud2-GFP localized at specific positions (bud neck or CRMs). Bud2-GFP in WT cells and *shs1-Δ1CTE* cells with or without Tm treatment. A large portion of Bud2-GFP was dispersed in ER stressed WT and ER stressed or unstressed *shs1-Δ1CTE* cells.

**Supplemental Figure 2: SPB duplication and positioning were monitored in WT, *slt2Δ1* and shs1-*Δ*1CTE cells using Spc110-GFP**

(A) Fields WT, *slt2Δ1*, and shs1-*Δ*1CTE cells showing SPB duplication and positioning visualized by a SPB reporter, SPC110-GFP, with or without Tm treatment.

**Supplemental Figure 3**

